# Accounting for spatial interactions in the upscaling of ecosystem services

**DOI:** 10.1101/2025.03.22.644705

**Authors:** Andrea Larissa Boesing, Gaëtane Le Provost, Margot Neyret, Anja Linstädter, Javier Muro, Jörg Müller, Kirsten Jung, Markus Fischer, Maximilian Lange, Olena Dubovyk, Paul Magdon, Ralph Bolliger, Sophia Leimer, Steffen Boch, Swen Renner, Till Kleinebecker, Ute Hamer, Valentin H. Klaus, Wolfgang Wilcke, Peter Manning

## Abstract

1. Maps of ecosystem service (ES) supply are frequently used to guide spatial planning, policy making, and ecosystem management. However, these are typically based upon coarse land-cover proxies. This approach lacks a strong mechanistic basis, and neglects spatial biodiversity dynamics and interactions among landscape properties that can modify ES provision.
2. We present an analytical framework for ES upscaling that incorporates spatial interactions between landscape properties to determine ES supply. The resulting models can be viewed as a spatially informed ES production function. The approach comprises seven steps that include several elements absent from most existing approaches, notably a procedure for identifying geodata variables that represent the true mechanistic drivers, the inclusion of spatial interactions in the upscaling model, and modification following expert feedback on the selected model.
3. We demonstrate the approach using two example ES from German grasslands: biodiversity conservation and water supply. We show that the inclusion of spatial interactions in the upscaling model improved model predictions from 15% to 33% depending on the ES evaluated. In addition, inclusion of spatial interactions led to reduced error associated with the upscaled estimates.
4. By overcoming several shortcomings of existing, upscaling approaches we generate resulting maps of ES supply that can more reliably inform spatial planning Further, the underlying models allow for simulation of changes in the drivers of ES supply and estimation of respective outcomes. These advantages have the potential to better link detailed local-scale ecological understanding and land management with large-scale ES supply mapping, and thus better inform decision making and spatial planning.

## 1. INTRODUCTION

For many different purposes within ecology and land management there is a need to upscale small-scale, often plot-level, ecological measurements to larger spatial scales at which most ecosystem management decisions are made (Maes *et al*. 2012). This is particularly true for ecosystem services (ES) which are mapped in both research projects and in national and regional assessments to inform environmental planning and communicate ecological information to stakeholders (Nelson *et al*. 2009; IPBES 2019).

Ecosystem service mapping is conducted using a range of methods. Perhaps the simplest and most widely applied is the use of land use proxies. Here, ES supply is derived from land cover maps, with each unit of land cover typically having a fixed ES supply rate across its entire area (McGuire *et al*. 2001; Nelson *et al*. 2009). This approach is often used in the absence of detailed field data, but can be inaccurate as many other factors such as abiotic factors, specific biodiversity components or landscape structure also drive ES supply (Eigenbrod *et al*. 2010). Errors associated with this approach can further propagate when ES valuation applies an estimate of economic value to a land unit based only on its land cover (Nelson *et al*. 2009; Plummer 2009; Bateman *et al*. 2013). While this simple approach is common significant progress in mapping and upscaling ES has been made by integrating more detailed biological data, such as vegetation indices or functional traits (e.g., Lavorel *et al*. 2011; Lavorel & Grigulis 2012), and incorporating a wide range of drivers operating at different spatial scales (Martínez-Harms *et al*. 2016; Spake *et al*. 2019; González-Chaves *et al*. 2022).

Recent years have also seen the emergence and rapid development of machine learning techniques for ES mapping, in which automated procedures select and generate high performing models from large number of candidate variables (Willcock *et al*. 2018; Scowen *et al*. 2021). Although machine learning techniques were designed to capture complex systems, the resulting models often lack a clear mechanistic/causal link to the service in question, mostly due to lack of a hypothesis-based selection of variables. This also leads to issues regarding the interpretability of the selected model and its predictions (Perry *et al*. 2022). A shift towards a more mechanistic representation of the relationships between ES and their drivers (van der Plas *et al*. 2019) should provide more reliable mapping, and allow for a better assessment of the causes of trade-offs and synergies between multiple ES (Neyret *et al*. 2021), resulting in better informed decision making.

An important aspect missing from many ES upscaling approaches is the incorporation of spatial processes and interactions between landscape components. While existing approaches discuss the integration of multiple scales of effects when upscaling ES (e.g., Spake *et al*. 2017, 2019) very few consider spatial interactions and processes explicitly (Le Clec’h *et al*. 2019; González-Chaves *et al*. 2022). This is potentially problematic as many landscape ecology studies have shown that ES involving the spatial movement of matter and organisms are strongly influenced by landscape composition and configuration, including spatial biodiversity patterns (Martin *et al*. 2019; Le Provost *et al*. 2023). Indeed, when spatial interactions have been accounted for in ES upscaling they are shown to be very important for multiple ES, with ES supply estimates changing by ∼ 20-700% (min-max) depending on the ES considered (Verhagen *et al*. 2016). Even though landscape ecology has produced exceptional knowledge in the past decades regarding the impact of landscape structure and spatial interactions driving ecological processes and biodiversity-ecosystem functioning relationships (Fahrig 2003; Tscharntke *et al*. 2012; Mitchell *et al*. 2015), this knowledge is still poorly integrated into ES upscaling and mapping approaches. For example, detailed information on how surrounding habitats affect crop pollinator supply (e.g., González-Chaves *et al*. 2020) is rarely upscaled beyond study areas (but see González-Chaves *et al*. 2022).

Spatially explicit and mechanistic upscaling of ES and their drivers would also be useful in biodiversity-ecosystem function and biodiversity-ES research. In these closely related fields, a mechanistic understanding of how biodiversity change affects the landscape level supply of ES is still largely controversial (Klaus *et al*. 2020; Freitag *et al*. 2022), despite increased recognition that larger-scale processes are major drivers of biodiversity-ecosystem functioning relationships (Van Der Plas *et al*. 2016; Mori *et al*. 2018; Le Provost *et al*. 2023). Importantly, there is a mismatch between the scale at which relationships between biodiversity and ecosystem function are usually determined and the scale at which most management actions and policy decisions are performed, such as protective legislation and payments for ES (Manning *et al*. 2019; Huber *et al*. 2022). This mismatch represents a barrier to effective communication of the importance of biodiversity and ES to wider society (Mace *et al*. 2012; Isbell *et al*. 2017), and overcoming it requires upscaling these respective relationships.

Here, we present an ES upscaling framework that is based on statistical approximations of ES production functions (Fig. 1) rather than proxies of ES supply. Our framework advances previous approaches by: i) systematically assessing and incorporating the role of drivers across multiple levels of abiotic and biotic organization to formulate a robust and mechanistic statistical ES production function, and ii) including spatial interactions with the surrounding environment into the ES production function. Our framework encompasses seven key steps. We first describe the rationale behind each step and then demonstrate the application of the framework with two ES examples: biodiversity conservation and water supply, identified as important ES by multiple stakeholders in the focal study region. We used data from the large-scale and long-term German Biodiversity Exploratories project (Fischer *et al*. 2010), https://www.biodiversity-exploratories.de/en) to train the models and upscale.

**Figure 1.**
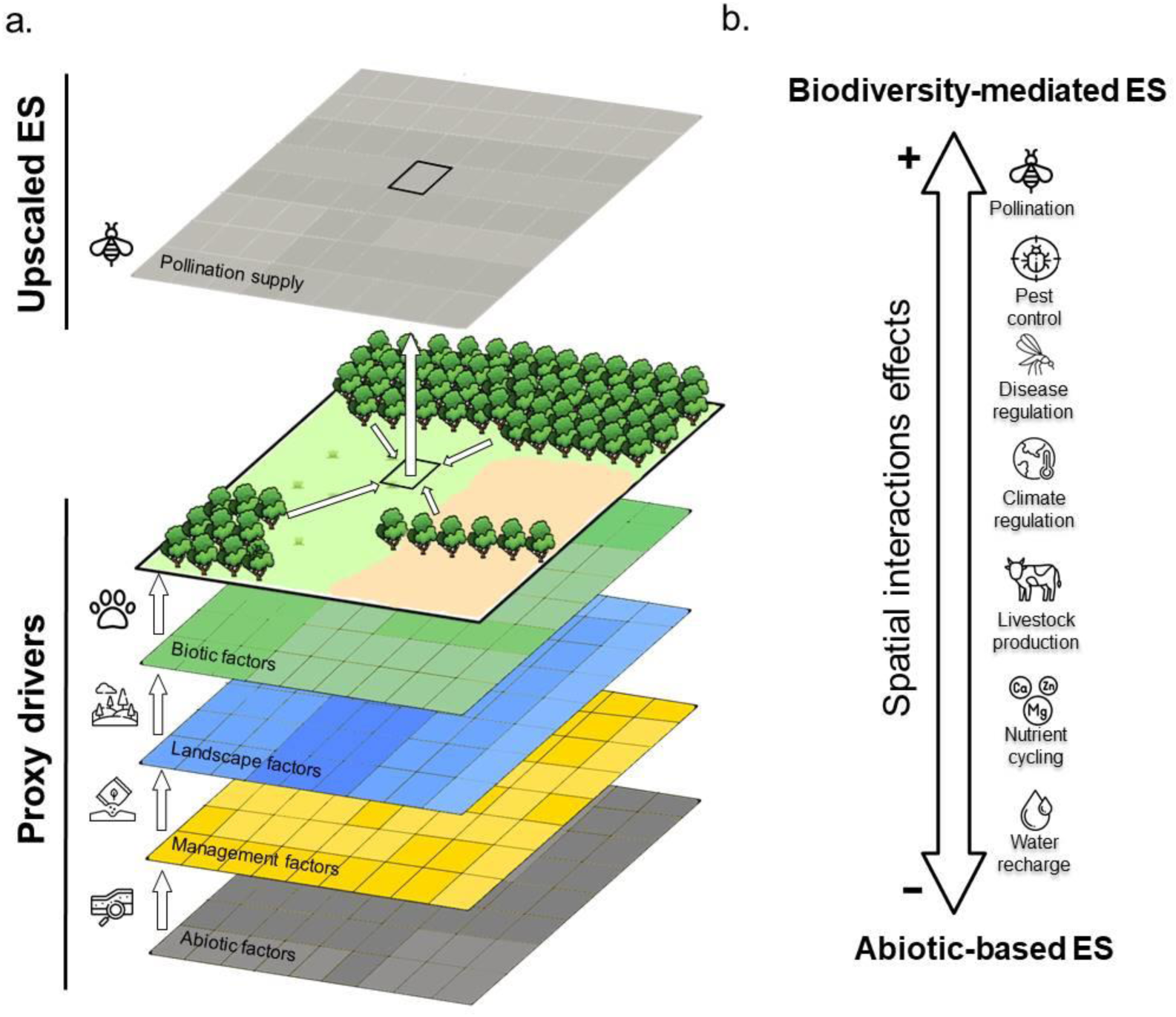
Representation of (a) spatially informed ecosystem service production function and (b) the expected importance of spatial interactions in driving the supply of different ES. The hierarchy of proxy drivers in (a) is abiotic factors, management factors, landscape factors, and biotic factors. We hypothesize that biodiversity-mediated ES (e.g., pollination, disease regulation) are typically more influenced by spatial interactions than ES dominated by abiotic factors (e.g., nutrient cycling, water recharge).

## 2. SEVEN STEPS TO UPSCALE ECOSYSTEM SERVICES

Our analytical framework comprises *seven* steps (Fig. 2). These are: 1. Define the area for which upscaling is required (*Define the area*). 2. Identify which ES are demanded by stakeholders and appropriate indicators for these (*Identify ecosystem services*). 3. Identify spatially explicit indicators of the potential biological and physical drivers of each ecosystem service (*Identify multiple proxy drivers*). 4. Select a semi-mechanistic and spatially explicit statistical model for upscaling (*Select the mechanistic model*). 5. Conduct expert evaluation of the validity of the selected model (*Expert evaluation*). 6. Adjust the models following expert advice and cross-validate the model to assess performance (*Adjust and validate*). 7. Upscale, acknowledging uncertainty in model predictions (*Upscale*) (Fig. 2).

**Figure 2.**
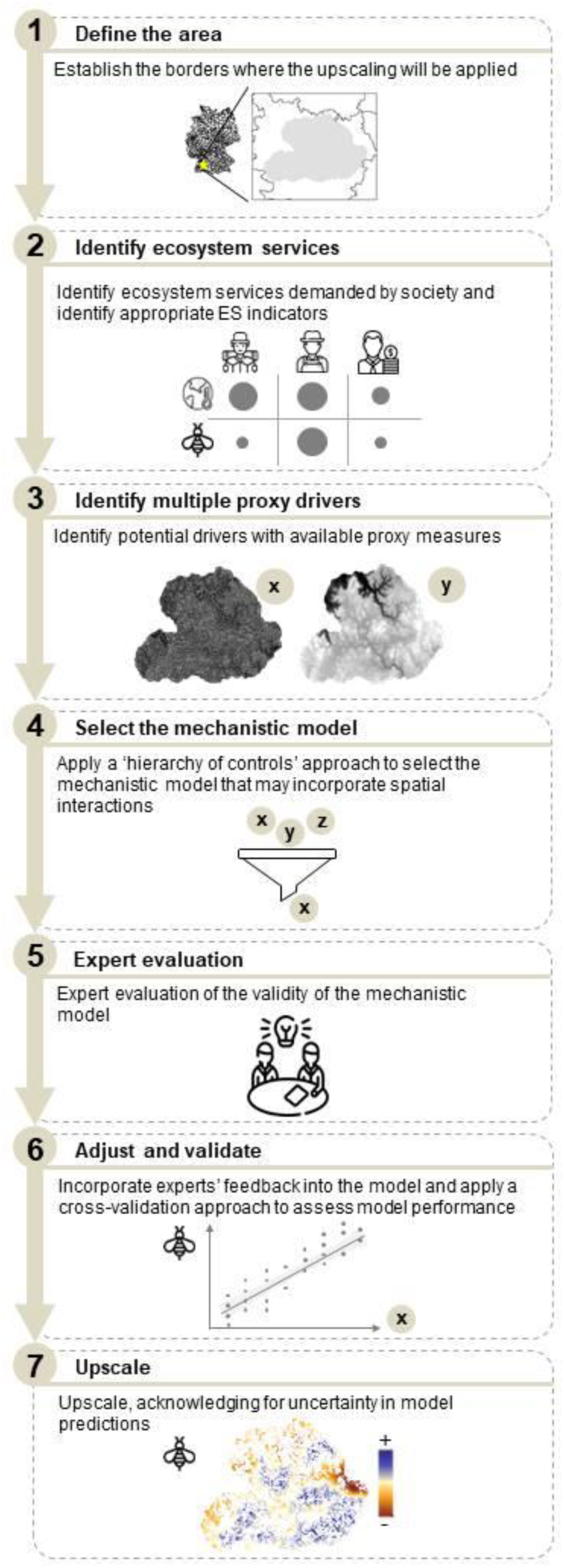
Seven-step analytical framework for spatially informed ecosystem service production function based upscaling. We present the procedures to be taken when high-quality plot-level data on ecosystem service supply and its drivers are available.

In the following sections we describe these steps in detail. The framework we present is likely to be most effective when detailed plot-level information relating to ES supply is available to inform upscaling, and so we concentrate on such cases. Such data is increasingly available (e.g., NutNet https://nutnet.org/; Biodiversity Exploratories https://www.biodiversity-exploratories.de/en/; FunDivEurope http://project.fundiveurope.eu/) but unfortunately, it is not available in many parts of the world.

### 2.1 Define the area

Upscaling is based on the transfer of values from one or more study sites to a larger geographical setting. In some cases, predictive models are applied outside the original study area to new geographic contexts that might differ from the training data in their environmental properties (Meyer & Pebesma 2021), potentially leading to erroneous predictions. Thus, before upscaling, the boundaries of the area to be predicted must be defined to ensure that appropriate indicators, proxies and baseline data are used. As ES by definition involve a human element, e.g., in terms of the demand and identity of ES, it may also be appropriate to define this boundary as that of the socio-ecological-system. Whilst ecological boundaries are usually defined by interfaces between different ecosystems or communities, social boundaries are typically set by institutional forces, and represented as cultural and/or administrative borders (Dallimer & Strange 2015; Ropero *et al*. 2021).

### 2.2. Identify ecosystem services

Ecosystem services are related to human demands, and it is therefore important to map the ES that are demanded by people and the regulating services that underpin these. Such a task requires a multidisciplinary approach involving social scientists to accurately assess stakeholder’s ES needs and preferences (Peter *et al*. 2022; Neyret *et al*. 2023).

Once the demanded ES are identified, it is necessary to identify appropriate *indicators* (Müller & Burkhard 2012) that capture the characteristics and status of a given ES, and which can be measured and upscaled appropriately (Layke *et al*. 2012). For instance, carbon storage and/or sequestration rates can be used as an indicator for large-scale climate regulation (Smith *et al*. 2013; Maes *et al*. 2016); pollinator abundance as an indicator of pollination (Woodcock *et al*. 2019), and the number of species in priority taxa can indicate biodiversity conservation value (Veach *et al*. 2017). Such indicators are usually only proxies of the ES supply and already introduce uncertainty in the modeling process. Thus, good indicators minimize uncertainty. ES indicators are ideally collected at the plot-level in a well-replicated study design which encompasses the full variability within the defined region (Redlich *et al*. 2022). Direct ES indicators should be preferred over indirect ones as the former more closely relate to mechanistic drivers of the final ES (Richter *et al*. 2021).

### 2.3 Identify multiple proxy drivers

Before creating a mechanistic statistical model for upscaling, it is necessary to identify multiple potential drivers of the ES indicator. The first step in this process is to review existing knowledge of the ES indicator, and select candidate variables that may drive the ES, including, for instance, abiotic drivers, land use, landscape features (e.g., surrounding land cover and its configuration) and biotic properties (e.g., species richness and composition). Next, we need to identify reliable *proxy* measures of these driver variables, e.g., remotely sensed measures or those mapped via conventional means. These should be available at the landscape scale whilst also accurately representing the true drivers. Advances in remote sensing data in recent years have substantially improved the resolution, accuracy, and temporal range of the available data e.g., to derive landscape structure drivers (Cavender-Bares *et al*. 2022).

We advocate the use of proxies from four classes of variables, which form a ‘hierarchy of controls’ (Fig.1 A). These are: abiotic factors, management factors, landscape factors, and biotic factors. *Abiotic factors* such as climate and soil attributes (soil type, soil pH, soil texture, soil depth, etc.) underpin many ES (e.g., nutrient cycling, soil carbon storage) and data are often available at regional and national levels (e.g., Copernicus, European Soil Data Centre). Topographic and geological properties, such as slope, aspect, and topographic wetness index, may also be important abiotic drivers and can derived from Digital Terrain Models (DTMs) and geological maps. *Management factors* such as management intensity and type are often important local-level predictors of ES, especially for provisioning services such as crop, livestock and fodder production. Proxy measures of management factors might be difficult to identify, but new techniques in remotely sensing anthropogenic activities are constantly improving, making reliable information describing forest disturbance, crop and grassland type and land use intensity available at large spatial scales (e.g., Griffiths *et al*. 2020; Lange *et al*. 2022; Schwieder *et al*. 2022). Additional data sources include agricultural census data and agricultural soil suitability maps (Le Clec’h *et al*. 2019). *Landscape factors* characterize the surroundings of the focal plots from which the ES are measured. They include landscape composition (i.e., land-cover types) and configuration (i.e., land-cover arrangement within a landscape), as well as land management measures at the landscape scale that can be derived from remote-sensing approaches (Griffiths *et al*. 2020; Lange *et al*. 2022). Finally, *biotic factors* include variables such as species richness and composition, biomass, and functional composition. The availability and quality of remote sensed data for biotic factors has dramatically improved in several domains, including plant species richness and composition (e.g., Civantos-Gómez *et al*. 2021; Muro *et al*. 2022), but also functional traits (Martínez *et al*. 2016). The examples cited here provide a flavor of the wide range of variables that can be employed, and which together form the main drivers of ecosystem service supply (Potschin & Haines-Young 2011).

The next step is to determine the ‘hierarchy of effects’ of the chosen proxy drivers, based upon the principles of hierarchy theory (Allen & Starr 1982; Müller 1992). Hierarchy theory posits that higher-level drivers constrain or moderate the effects of lower-level drivers, which can interact, in turn, to influence higher-level outcomes (Allen & Starr 1982). Lower hierarchy is influenced by the upper-level hierarchy, but in most of the cases, not vice versa (Fig. 1A) In addition, higher levels generally change more slowly than lower levels, and very often, the spatial extent of higher-level factors is broader than that of lower levels (Müller 1992). For instance, the topography of a region (which varies at larger scales) may influence management practices at smaller scales (e.g., mowing or fertilization regimes), but the opposite is rarely true since these lower-level drivers are inherent properties (Dominati *et al*. 2010). Similarly, fundamental abiotic conditions such as climate, soil pH, soil type and texture affect biodiversity (Le Provost *et al*. 2021), but the opposite is also unlikely to happen (although there are exceptions). The hierarchy of controls approach has been successfully applied in a range of studies aiming to predict ecosystem functions from biotic factors (e.g., Díaz *et al*. 2007; Lavorel *et al*. 2011; Manning *et al*. 2015). Moreover, modeling ES supply as driven by a hierarchy of controls is also consistent with the widely used ES cascade model (Potschin & Haines-Young 2011; Zhang *et al*. 2022), in which abiotic features alter the biota and together with external inputs produce ES supply (Potschin & Haines-Young 2011; Bateman & Mace 2020).

While the supply of some ES at any given point in space is primarily driven by factors that act locally (e.g., soil carbon storage or livestock production; Le Provost *et al*. 2023), other ES, especially those associated with mobile organisms, such as pollination, are also regulated by factors operating at the landscape scale (Le Clec’h *et al*. 2019; Le Provost *et al*. 2023). For such drivers it is paramount to determine the *scale of effect*, i.e., the scale over which the process affecting the ES operates and so best approximates the observed pattern (Jackson & Fahrig 2015). Since the scale of effect of a given driver is often unknown, a multi-scale approach should be performed. Here, variables incorporating a range of scales are used in modeling (e.g., land cover measures in radii of 200, 500 and 1000m) and the ‘best’ is selected as an approximation of the scale of influence that a driver operates over (Jackson & Fahrig 2015). Selection of the scale of effect can be based on the strength of the relationship between the landscape property and the ES indicator (see Jackson & Fahrig 2015).

### 2.4 Select the mechanistic model

In the next step of our upscaling framework, we employ the hierarchy of controls approach to a statistical model selection approach. In this, proxy drivers are fitted in a fixed sequential order to identify and select the true mechanistic drivers of ES supply. In practical terms, groups of terms are added to a regression model in fixed sequential order, determined by their influence across levels. The order of hierarchy is such that variables added later in the modeling process are unlikely influence the previous ones. At each step all potential driver variables of a given level of the hierarchy are added and their impact on model likelihood is assessed. Only terms found to significantly improve model likelihood in a likelihood ratio deletion test (LRTs) are retained. After conducting this procedure for all variables of a hierarchical level, an interim model is selected, and the modeling process moves to the next level of the hierarchy, where another set of variables is added and the process is repeated (Fig. 3). This continues until all levels are added and the final model is selected. Interactions between variables within and across levels (see Spake *et al*. 2019) should be tested, since many indicators are driven by multiple drivers and their interactions. To avoid the retention of many weakly significant factors, we suggest using only lower level interactions.

**Figure 3.**
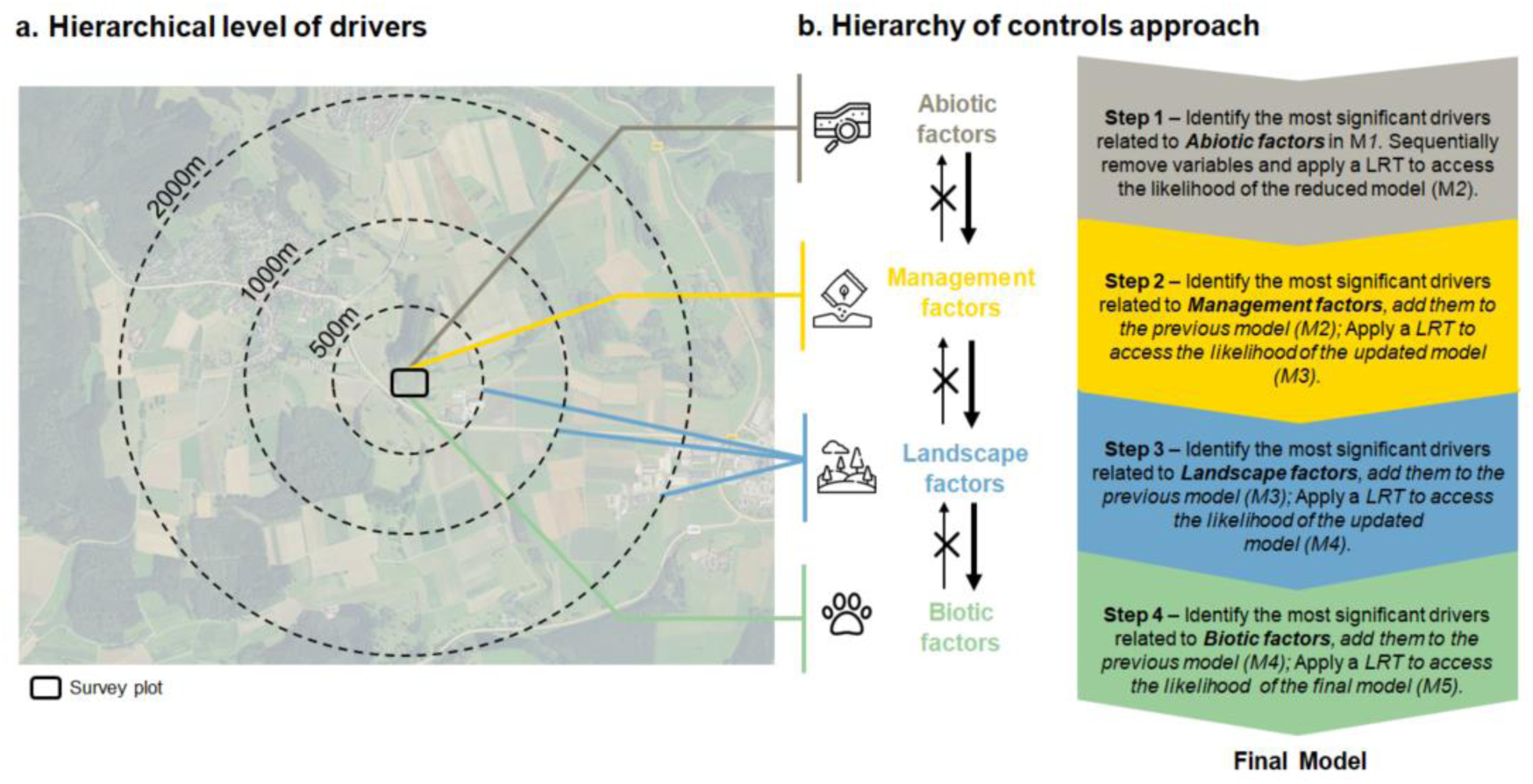
Representation of (a) the effects of proxy drivers across scales, and (b) the steps used in the hierarchy of controls approach to select the final mechanistic upscaling model.

After selecting the mechanistic upscaling model, we suggest checking whether spatial autocorrelation, and related issues of pseudo-replication, impacts the upscaling (Dormann 2007; Dormann *et al*. 2007). Spatial autocorrelation (i.e., the higher similarity of closer samples) is a common phenomenon in ecology that can appear for several reasons, including biotic processes such as dispersal (Miller 2012) or abiotic factors, such as climate, soils, topography and disturbances (Ranta *et al*. 1997). When spatial autocorrelation is related to the driver of interest, the inclusion of the variable causing the autocorrelation as fixed factor in the model may be sufficient. In other cases, ignoring spatial autocorrelation can lead to biased parameter estimates and artificially low standard errors (Albert & McShane 1995; Keitt *et al*. 2002). Detection of autocorrelation can be done in several ways including Moran’s I correlograms, Geary’s correlograms and semi-variograms (Dorman *et al*. 2007). Where spatial autocorrelation is detected, we suggest using autocovariate regression and spatial eigenvector mapping to capture the spatial configuration of covariates and adding a term for this to the upscaling model.

### 2.5 Expert evaluation

After determining the mechanistic model, we suggest consulting experts of the ES of interest for model evaluation. Here they should check if the driver variables included are likely to be true mechanistic drivers, or simply correlates of them. This step is important when dealing with multiple ES, as it is unlikely that the person performing upscaling will have expert knowledge of all relevant ES. Experts should also assess whether any other important drivers are missing, if the database used for the calculations is suitable, e.g. in terms of spatial and or temporal resolution, and if the selected variables represent the true mechanistic drivers the ES. After consultation, models must be revised and adapted, e.g., by substituting spurious drivers with closely correlated variables that better represent the true drivers. While this step adds complexity to the process, and costs time, model outputs are likely to be greatly improved, especially where the model is used to make predictions.

### 2.6 Adjust and validate

After expert assessment and revision, the next step is to assess model performance via cross-validation. The aim here is to estimate the accuracy of model predictions. In practice, we suggest evaluation of model performance and associated error by setting aside a random set of test observations at the model calibration stage and using these to quantify prediction error (true validation step). Validation techniques include ‘validation set aside’, ‘k-fold cross validation’ and ‘repeated k-fold cross-validation’ (Browne 2000), each of which has advantages and drawbacks. One limitation of cross-validation is that it is only possible for large datasets. For smaller datasets we recommend (repeated) k-fold cross-validation to estimate the prediction error rate as this can use the entire dataset for generating estimates. Knowing data limitations is crucial when looking for an error estimate, which in turn is important when communicating uncertainty. A model should be considered reliable when it retains the true drivers of ES change and also has low error associated with its estimates.

Finally, regarding the statistical setup of the models, both linear and generalized linear models may be used. Care must be taken when upscaling non-linear models and for the presence of outliers in the training data or in the proxy drivers as these can result in over or under-estimation of the service or result in higher errors. When outliers are present, one simple solution is to limit predictions to areas within the range of the training data. This may reduce the coverage of predictions but prevents extrapolation into areas where error may be high.

### 2.7 Upscale

Once the final model is selected and its performance assessed, the layers containing the drivers must be prepared and the landscape interactions incorporated when necessary. For non-spatial drivers the model is applied to each pixel of the driver raster, while for spatially interacting drivers a calculation needs to be made prior to upscaling of the ‘landscape surroundings’ values of each pixel (e.g., the proportion of area of a certain area habitat type within a certain radius of the pixel). The model is then applied to this layer of calculated values and the other proxy drivers for each point in the landscape to generate estimates (Fig. 1). Creating the spatially-explicit layers may be time consuming and computationally demanding, so we advise the integrated use of different tools, including QGIS, ArcGIS and R. The resulting maps will represent ES supply, as function of the mechanistic drivers.

## 3. Applying the upscaling framework in real landscapes

Our approach to upscale ES incorporates some novelties that will likely not only improve the ES predictions, but also improve their use as a tool for decision making. The integration of social aspects to define the needs of a region, the hierarchical approach to identify the mechanistic drivers of ES, and the integration of spatial processes in the upscaling makes this approach powerful.

Next, we demonstrate how our upscaling framework can be applied in practice, using two case studies: biodiversity conservation and water supply, both within the Schwäbische Alb region in southwest Germany. We performed the upscaling only for grassland areas, since the plot-level indicators were collected in grasslands and predictions outside this land use type would be unreliable.

### Step 1 – Define the area

In this case the study area was defined by a fundamental research project, the large-scale and long-term Biodiversity Exploratories project (Fischer *et al*. 2010), which was established to investigate land-use-biodiversity-ecosystem function relationships. Plots were selected to span the full range of land use intensity of each region. Recent work has extended plot-level work into the social-ecological systems surrounding the original study plots (Peter *et al*. 2022; Neyret *et al*. 2023), and the boundary of upscaling was defined by this broader project.

### Step 2 – Identify ecosystem services

Social surveys conducted with 116 respondents from 14 stakeholder groups identify 12 highly prioritized ES in the area (see Peter *et al*. 2022), including *biodiversity conservation* To represent biodiversity conservation, we used above-ground multidiversity, a standardized measure of diversity across multiple taxa as indicator (Allan *et al*. 2014). We used 14 above-ground taxa (Table S1) across 50 grassland plots sampled between 2008 and 2021. In our case, above-ground multidiversity was used as an indicator since multitrophic diversity is critical for sustaining ecosystem functioning (Soliveres *et al*. 2016).

As a second example, we use groundwater recharge, an indicator of water supply to people, crop, fodder and timber plants (also identified as important ES for the region). Values were derived from a soil water balance model developed to calculate vertical soil water fluxes from the top layer (0-0.15 m) (Kreutziger 2006; Leimer *et al*. 2014, 2018) (SM1). As input data for the model, we used biweekly rainfall data from 2010- 2016 in the 50 grassland plots and net water fluxes from soil to deeper layers were aggregated to annual values. We used the average across the seven years as indicator of water supply.

### Step 3 – Identify multiple proxy drivers

We collected data for a variety of proxy drivers that potentially influence the supply of these ES. These were organized in a hierarchy of: 1. abiotic factors (topographic variables such as slope, elevation, topographic wetness index, and aspect, and soil attributes, such as soil texture, pH, clay content, bulk density, soil depth), 2. management factors (including land use intensity (LUI; Blüthgen *et al*. 2012), grazing, fertilization and mowing at the local level), 3. landscape factors (500-2000 m scale; including both landscape composition and configuration of land-uses, especially woodlands, grasslands, and crops, and also patterns of landscape-management, as represented by amount and configuration of low and high LUI grasslands) at the best scale of response (1500m surrounding the plots for biodiversity conservation, and 2000m for water recharge), and biotic factors (as mean alpha plant richness). Details about the mechanisms behind these drivers, metrics, scales and sources of remote sensing data for the proxy drivers can be found in Supplementary Material 2. Correlations across variables were checked and variables with r =<0.7 were used.

### Step 4 – Select the mechanistic model

We next applied the ‘hierarchy of controls’ approach to identify mechanistic upscaling models. For each hierarchical level (Fig. 3), we tested which variables significantly improve model likelihood in a likelihood ratio deletion test (LRT), before advancing to the next step of the modeling process. For both biodiversity conservation and water supply, we used linear models with a Gaussian distribution. Model fit was evaluated by examining the residuals of the relationship (Hartig 2020) and we tested for spatial autocorrelation using Moran’s I correlograms (Paradis & Schliep 2019). The initially selected model for biodiversity conservation explained 37% of variation and contained terms related to abiotic factors (phosphorus content at local level), landscape factors (landscape heterogeneity, measured as the diversity of land use types at 1500m scale), and biotic factors (estimated plant richness, at local level). For water supply, the initial model explained 72% of the variance and contained terms for abiotic factors (pH and bulk density at local level), and landscape factors (% of coniferous cover at 2000m scale, and % high LUI cover, 1500m scale).

### Step 5 – Expert evaluation

We sent the initially selected models to experts from the Biodiversity Exploratories Consortium who were the original data collectors of the ES indicators in the form of a spreadsheet containing the models, and requested feedback in terms of model plausibility, such as appropriateness of driver variables selected, absence of key drivers and direction of effects. For *biodiversity conservation*, three changes were made following this consultancy: first, we updated landscape LUI measures as improved data had become available (Lange *et al*. 2022); second, experts pointed out potential circularity in using remote sensed plant richness estimates as proxy driver of biodiversity conservation since related measures were also included in the multi-diversity index, thus it was removed; finally, we moved topography measures to the first level of hierarchy, before soil attributes, in the hierarchy of controls approach. For water supply, two changes were also performed: the model initially contained bulk density and LUI at the landscape (1500m) scale as drivers, but the direction of effect was identified as spurious by experts. Second, we moved topography measures to the first level of hierarchy.

### Step 6 – Adjust and validate

Models were re-fitted following expert feedback. The final model after revision for *biodiversity conservation* had a R^2^ of 0.47, and contained additive effects of land-use diversity at the landscape scale (1500m) (explaining 15% of variance), slope (14%), soil depth (10%), and soil phosphorus content (8%) (Table S2). The presence of a landscape factor in the model indicates the importance of spatial processes in driving biodiversity, consistent with biological studies of the same grasslands (Le Provost *et al*. 2021). The final mechanistic upscaling model for *water supply* had a R^2^ of 0.56, and contained additive effects of soil type (explaining 26% of the variance) and the percentage of coniferous forest (at 2000m scale, 25% of variance) (Table S2). These factors influence water permeability and thus infiltration rates; for instance, coniferous forests intercept more water than broad-leaved and deciduous forests, and the effect on water infiltration reduction is influenced by the density of plantations (Bellot *et al*. 1999; Adane & Gates 2015). See SM2 for a detailed description of the mechanisms driving the supply of both ES.

We used a K-cross fold validation to access model predictability power, where the entire dataset is used to calculate the root mean squared error (RSME) for both models, since our training dataset is relatively small (N=50). In general, lower RSME, represents lower prediction error, and it indicates the degree of correspondence between observed and predicted values. For *biodiversity conservation*, we obtained a RSME of 0.08 (observed: 0.13-0.58, predicted: 0.0001-0.71). Given the range of both training data and predicted values, a RSME of 0.08 could be considered acceptable error (i.e., that the predictions may vary ± 0.07 from the actual values). For *water supply*, we obtained a RSME of 53.11 (observed: 477.69-791.51mm, predicted mm: 339.39-737.39mm).

### Step 7 – Upscale

Remote sensed inputs were harmonized and stacked together, and the model coefficients were applied to each pixel of the stack to make predictions (details in the Tutorial). First, we illustrate how spatial interactions, in the form of the land use diversity surrounding grasslands affect supply levels of *biodiversity conservation*. The model predicts that grasslands surrounded by a high-diversity of land-use types, at steeper slopes, and where the soil is both deep and poor in phosphorus content, possess the highest biodiversity conservation value.

Second, for water supply, areas embedded in landscapes with lower coniferous cover, and with soil of type Braunerde-Terra Fusca, have a higher capability to increase groundwater recharge in our study system (Fig. 5).

**Figure 4.**
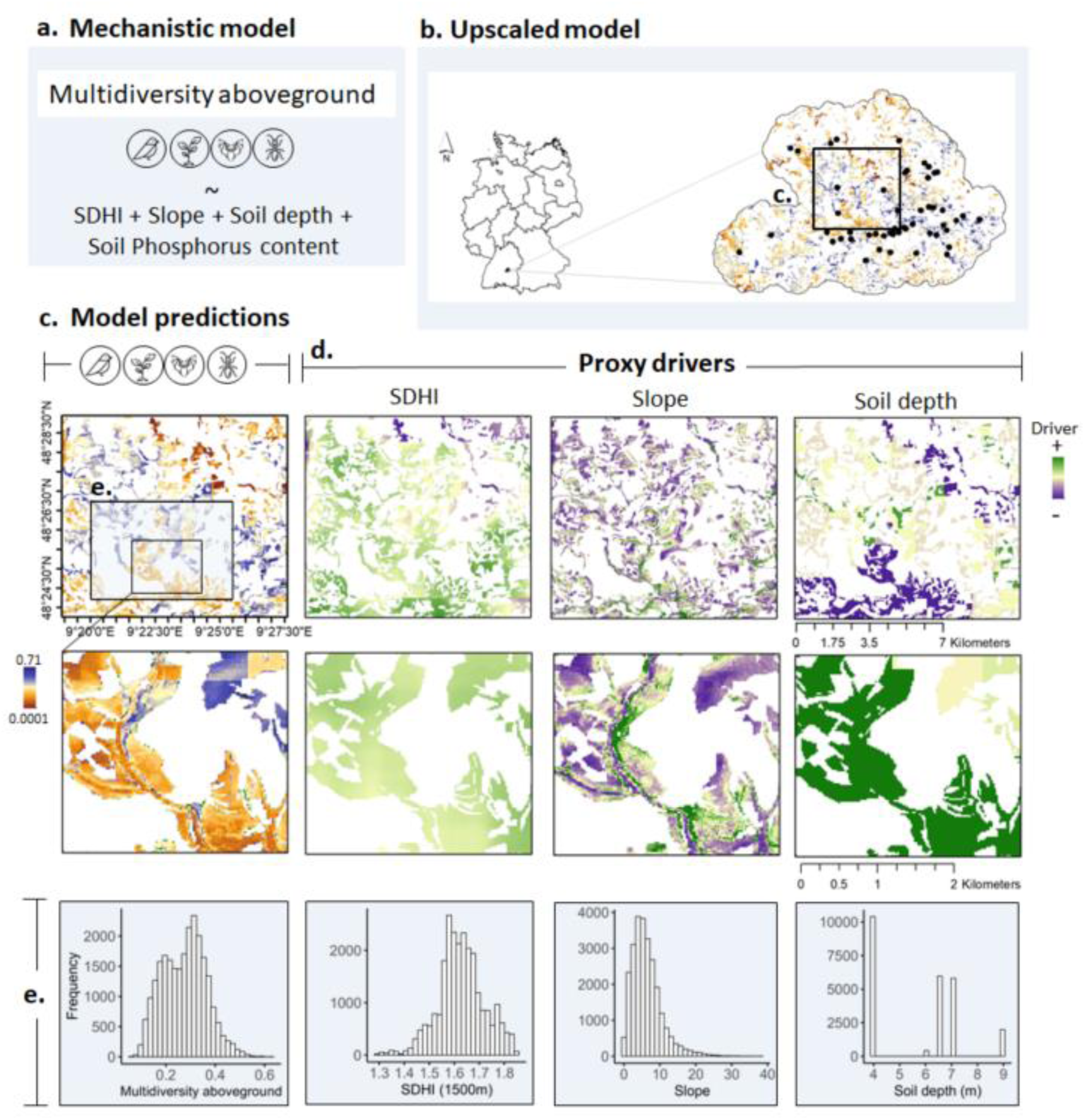
Upscaled predictions for biodiversity conservation (using multidiversity aboveground as indicator) for the studied region. We present the final upscaling model in (a). In (b) we present the region and the upscaled biodiversity conservation with a zoom in (c). The dots in (b) represent the plot locations from where the data was used to train the model. (d) presents three of the drivers used to model multidiversity aboveground. The middle row shows a closer zoom to show how spatial context, in the case of landscape heterogeneity (SDHI; Shannon’s diversity index) alters the supply of biodiversity conservation. The additive effect of SDHI and local factors (slope, soil depth, and soil phosphorus content) are noticeable, and landscapes with higher diversity of land uses have to higher multidiversity aboveground, and thus higher biodiversity conservation value. The bottom row (e) shows the data extract from the closer zoom (larger square), regarding the predicted values of multidiversity aboveground and respective predictor drivers (SDHI, slope and soil depth). Soil phosphorus content is not shown in the figure but included in the model. White areas in the map are areas outside the range of predictions.

**Figure 5.**
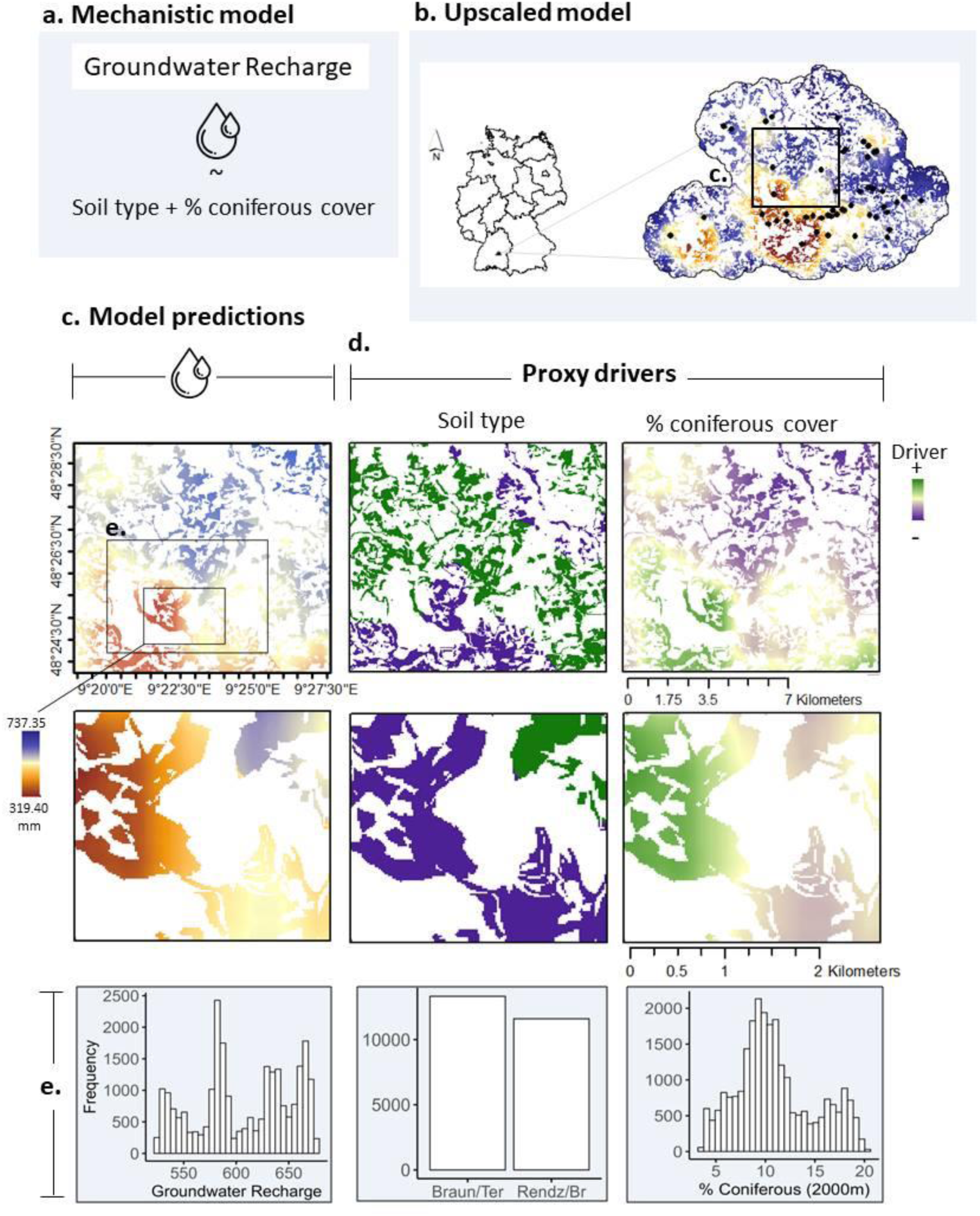
Upscaled predictions for water supply (using groundwater recharge as indicator) for the studied region. We present the final upscaling model in (a). In (b) we present the region and the upscaled water supply with a zoom in (c). The dots in (b) represent the plot locations from where the data was used to train the model. In (d) are presented the drivers used to model groundwater recharge (soil type and coniferous cover). The middle row shows a closer zoom within the smaller square to show how spatial context, in particular the cover of coniferous forest, alters groundwater recharge. Areas embedded in landscapes with lower coniferous cover, and with soil of Braunerde-Terra Fusca type, have a higher groundwater recharge and thus higher water supply. The bottom row (e) shows the data extract from the closer zoom (larger square), regarding the predicted values of groundwater recharge and respective predictor drivers (soil type and coniferous cover). White areas in the map are outside the range of predictions.

## 4. UPSCALING WITH VS WITHOUT SPATIAL INTERACTIONS

Integrating spatial interactions should provide more reliable estimates of ES supply if these processes are important, and we test this idea in thew following section. Since the landscape aspects are added as one of the last steps of the hierarchy of controls (before biotic factors), we did this by comparing the best models that either did or did not contain spatial interaction terms. For *biodiversity conservation*, removing landscape heterogeneity from the final mechanistic upscaling model, reduced explained variance from 47% to 32% and increased the RSME from 0.07 to 0.09. In the absence of the spatial drivers, the model underestimated multidiversity in more heterogeneous landscapes, and overestimated multidiversity in more homogenous landscapes (Fig. 6a,b).

**Figure 6.**
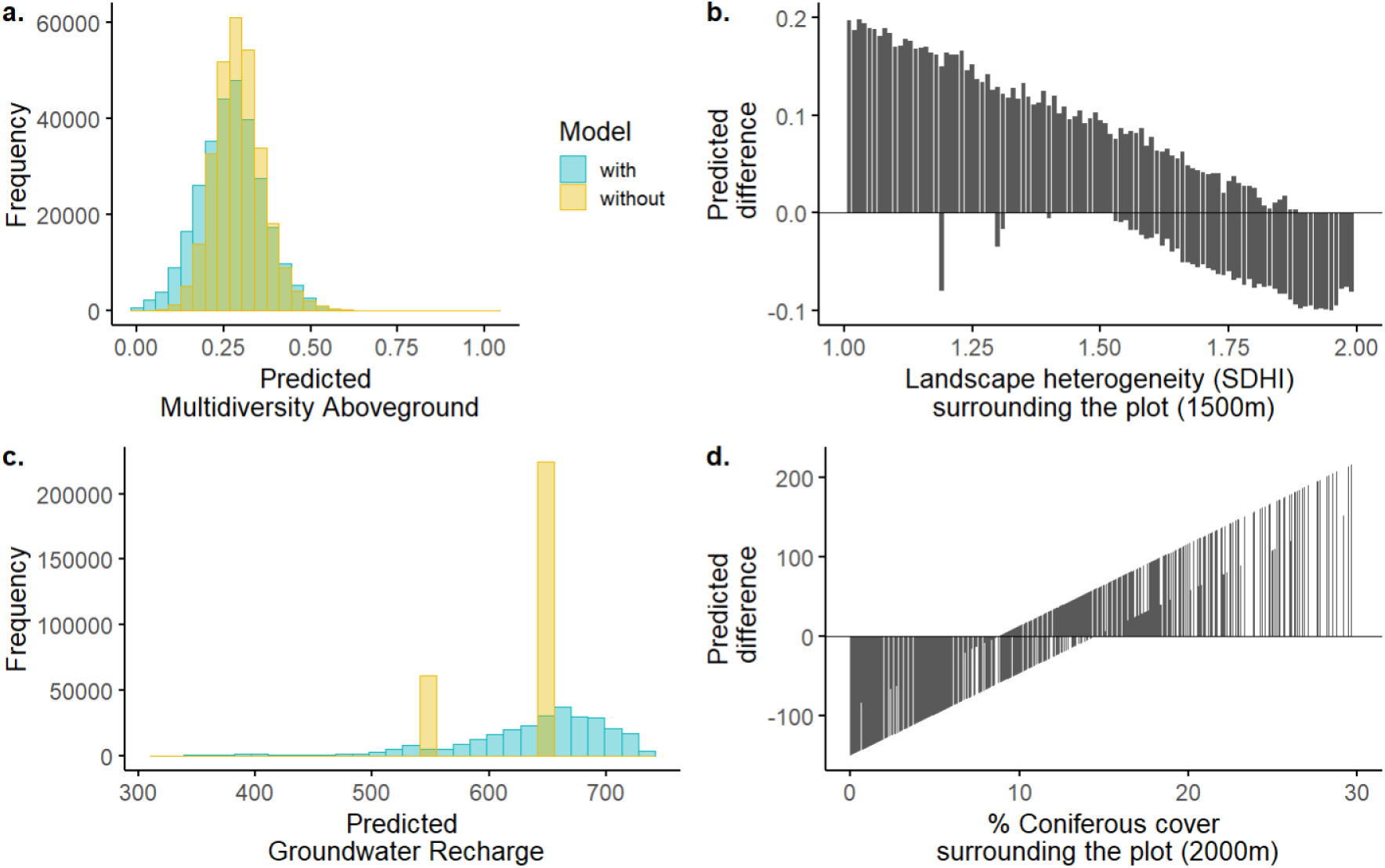
(a) Overall predicted estimates of multidiversity aboveground (biodiversity conservation) including spatial and non-spatial aspects (in this case, landscape heterogeneity measured by the Shannon’s diversity index, SDHI); (b) difference (without spatial interactions – with spatial interactions) on predicted estimates of multidiversity aboveground along the landscape change gradient; (c) overall predicted estimates of groundwater recharge (water supply) including spatial and non-spatial aspects (in this case, % of coniferous cover); and (d) difference (without spatial interactions – with spatial interactions) on predicted estimates along the gradient of coniferous cover.

For *water supply*, removing the landscape component reduced the explained variance by 25% and increased the RSME from 53.11 to 67.07. In the absence of spatial drivers, the model underestimates groundwater recharge by up to ∼100mm at lower coniferous cover (Fig. 6c,d), and overestimates it in landscapes with higher coniferous cover, by as much as ∼200mm in areas up to 30% of cover (Fig. 6c,d). Considering the mean of 626mm in the original data, such difference might correspond to around 32% in the estimations. Spatial model predictions ranged from 319.39 – 739.35mm, while in the non-spatial model only two values are predicted, based on coarse soil type data (646.41mm in Braunerde-Terra fusca soil type, and 549.80mm in Rendzina-Braunerde (Fig. 6c). These examples demonstrate that incorporating spatial interactions can strongly improve model performance and that these improvements are sufficiently great to impact and improve ES maps for decision making.

## 5. CONCLUSION

Our upscaling framework advances upon existing ES mapping approaches by modeling ES in mechanistic models that can incorporate spatial interactions between landscape components. The resulting maps have the potential to more reliably inform policy decisions, and spatial planning, for example in prioritizing areas for grants or subsidies to support ES provision, or spatial targeting of monitoring schemes. Further, as the resulting models are mechanistic, any change in the drivers can be simulated and its impact on ES supply estimated. For instance, in the case of our examples, the diversity of land-use types could be changed within a GIS model to assess the impact on the ES biodiversity conservation, and changes in coniferous forest cover could also be simulated to predict changes in groundwater recharge.

If widely applied, the proposed approach may form a bridge between detailed local scale ecological studies in which mechanisms are well understood, and larger-scale ecosystems service research, which tends to be performed at scales relevant to policy and planning. In particular, it allows assessment and quantification of the impact of compositional and configurational changes within landscapes at larger scales than they are usually assessed in landscape ecology research.

## ACKNOWLEDGEMENTS

We thank the managers of the Biodiversity Exploratories for their work in maintaining the plot and project infrastructure and for support via the central office. Markus Fischer, Eduard Linsenmair, Dominik Hessenmöller, Daniel Prati, Ingo Schöning, François Buscot, Ernst-Detlef Schulze, Wolfgang W. Weisser, and Elisabeth Kalko helped establish the Biodiversity Exploratories project. We thank the administration of the UNESCO Biosphere Reserve Swabian Alb as well as all landowners for collaboration.We also thank Eric Allan for supporting this manuscript with both tools and data. The work has been (partly) funded by the DFG Priority Program 1374 “BiodiversityExploratories”, specifically the BEF-Up II DFG contributing project (MA 7144/1-2). AL and OD acknowledge funding via the SEBAS DFG contributing project (LI 1842/4–1 and DU 1596/1–1). VK acknowledges funding by the Agroscope research program *Indicate*.

## AUTHORS CONTRIBUTIONS

A.L.B & P.M. conceived the idea and methodology. A.L.B. analyzed the data and led the manuscript writing; K.J., M.L., S.L., S.B., T.K., U.H., V.H.K., W.W. contributed discussing the models; A.L., J.M., J.M., M.F., M.L., O.D., P.M.,R.B.,S.L.,S.B., S.R.,T.K.,W.W. contributed with data. All authors commented upon the paper and contributed discussing the models.

## DATA AVAILABILITY

This work is based on data from projects of the Biodiversity Exploratories programme (DFG Priority Program 1374). Most datasets are publicly available in the Biodiversity Exploratories Information System (http://doi.org/10.17616/R32P9Q). IDs:27707,27706,21707,31442,31444,31371,31214,27087,27729,27727,31234,11541). Code used to prepare data and conduct analyses will be made available in the same repositoryAny other relevant data are available from the corresponding author upon reasonable request.

## Supplementary Material

### S1 - ES indicators used to measure Biodiversity Conservation and Groundwater recharge

Here we present the details of the indicators used to measure both Biodiversity conservation (using multidiversity aboveground as indicator) and Groundwater recharge.

#### Biodiversity conservation

We used the aboveground multidiversity (a standardized measure of the diversity across multiple taxa; such index may identify areas of high conservation priority (Allan *et al*. 2014)), thus a good indicator of Biodiversity conservation. The multidiversity index operates similarly to indices of multifunctionality used in BEF research (Kleijn *et al*. 2012) and to the World Wildlife Fund’s living planet, which quantifies the overall state of biodiversity (Loh *et al*. 2005). We used 14 above-ground taxa to calculate the Multidiversity above-ground (Allan *et al*. 2014) across 50 grassland plots collected between 2008 and 2021.

Multidiversity was calculated using the Multidiversity index (Allan *et al*. 2014) using no threshold parameter. In our analysis, we assumed that all of the taxonomic groups were of equal importance and calculated multidiversity as the average proportional species richness across taxonomic groups. We used14 aboveground groups (Table S1): Aranea, bats, Coleoptera, Dermaptera, Dictyoptera, Hemiptera, Hymenoptera, lichens, bryophytes, Neuroptera, Opiliones, Orthoptera, vascular plants and birds. Data is available in the databases described below and the code available in Tutorial 1.

#### Groundwater recharge

We used a soil water balance model, developed to calculate vertical soil water fluxes (in mm = L m^−2^) from one soil layer (0-0.15 m, which was selected because soils in the Schwäbische Alb are shallow and frequently do not reach deeper than 0.15 m) in grassland (Kreutziger 2006; Leimer *et al*. 2014, 2018) based on DVWK (1996). We used the net flux, which is the downward waterflux minus upward waterflux (capillary rise). As input data for the model, we used a biweekly rainfall dataset for each of the 50 grassland EPs, derived from rainfall measurements by the BECycles project and data from the Deutscher Wetterdienst (DWD), and climate data (soil moisture, air temperature, relative humidity) from the climate stations on the plots from 2010-2016. The measure unit is the water flux into deeper soil layers per year (mm). Data is available on Bexis 22746 and 27087 (Table S1).

**Table S1.** Datasets and their respective id number available in the Biodiversity Exploratories database (https://www.bexis.uni-jena.de/) used to calculate Multidiversity aboveground as indicator of biodiversity conservation and groundwater recharge. The datasets below were extracted from the synthesis dataset 27707; 27706)

| Dataset ID | Taxonomic group | Project in the Biodiversity Exploratories | Date (start) | Date (final) |
| --- | --- | --- | --- | --- |
| **Biodiversity conservation** |  |  |  |  |
| 27706 | Species information | Synthesis | 1/1/2008 | 31/12/2020 |
| 27707 | raw diversity | Synthesis | 1/1/2008 | 31/12/2020 |
| 4140 | Moss | Botany (core) | 1/5/2007 | 1/8/2008 |
| 5522 | Lichen | Botany (core) | 1/6/2007 | 17/06/2009 |
| 16871 | Coleoptera | Arthropods I (core) | 1/6/2008 | 1/9/2008 |
| 16893 | Hemiptera | Arthropods I (core) | 1/6/2008 | 1/9/2008 |
| 16894 | Aranea | Arthropods I (core) | 1/6/2008 | 1/9/2008 |
| 16895 | Orthoptera/Dermatoptera/Dictoptera | Arthropods I (core) | 1/6/2008 | 1/9/2008 |
| 16896 | Opiliones | Arthropods I (core) | 1/6/2008 | 1/9/2008 |
| 16897 | Neuroptera | Arthropods I (core) | 1/6/2008 | 1/9/2008 |
| 16908 | Hymenoptera | Arthropods I (core) | 1/6/2008 | 1/9/2008 |
| 19849 | bats | Vertebrates I (core) | 1/6/2009 | 30/09/2009 |
| 21446 | birds | Vertebrates II (core) | 15/03/2008 | 15/06/2008 |
| 21447 | birds | Vertebrates II (core) | 15/03/2009 | 15/06/2009 |
| 21448 | birds | Vertebrates II (core) | 15/03/2010 | 15/06/2010 |
| 21449 | birds | Vertebrates II (core) | 15/03/2011 | 15/06/2011 |
| 24690 | birds | Vertebrates II (core) | 15/03/2012 | 15/06/2012 |
| 25306 | birds | LMT (core) | 15/03/2018 | 13/06/2018 |
| 27386 | plants | Botany (core) | 1/1/2008 | 31/12/2020 |
| **Groundwater recharge** |  |  |  |  |
| 22746 | water fluxes | BECycles | 1/1/2010 | 31/12/2016 |
| 27087 | ecosystem functions | Synthesis | 1/1/2008 | 31/12/2018 |

## SM 2 -Mechanistic drivers of Ecosystem Services supply (Biodiversity conservation and Groundwater recharge)

We collected data for a large variety of proxy drivers that could influence the supply of the studied services (biodiversity conservation and groundwater recharge) (Table S3,S4). These were organized in a straightforward hierarchy of: 1. abiotic factors (topographic variables such as slope, elevation, topographic wetness index (TWI), and aspect, and soil attributes, such as soil texture, pH, clay content, soil type, and, soil depth), 2. management factors (including land use intensity (LUI), grazing, fertilization and mowing at the local level), 3. landscape factors (including both landscape composition and configuration of land-uses, including woodlands, grasslands, and crops, and also patterns of landscape-management, as represented by amount and configuration of low and high LUI grasslands), and biotic factors (as mean alpha plant richness) (Fig. S1). See Table S3 for details on drivers and sources and Table S4 for the proxy data tested in the models.

For the landscape factors, before including them in the hierarchy of controls selection, we tested each variable (e.g. grassland cover, grasslands edge density) at multiple spatial scales (500, 1000, 1500, 2000m) in order to select the scale of effect (Jackson & Fahrig 2015) and used the one with higher power of explanation (higher r^2^) in our models. We also checked for correlations across variables, and included only for uncorrelated ones. For Biodiversity conservation, the most significant landscape metric selected was the heterogeneity of land use types at 1500m, while for Groundwater recharge, both the % of coniferous forest (at 2000m) and the % of high-LUI cover (also at 2000m) were selected. However, we needed to leave LUI at the landscape scale out of the model, since this variable was negatively impacting model fit and model predictability.

**Figure S2.**
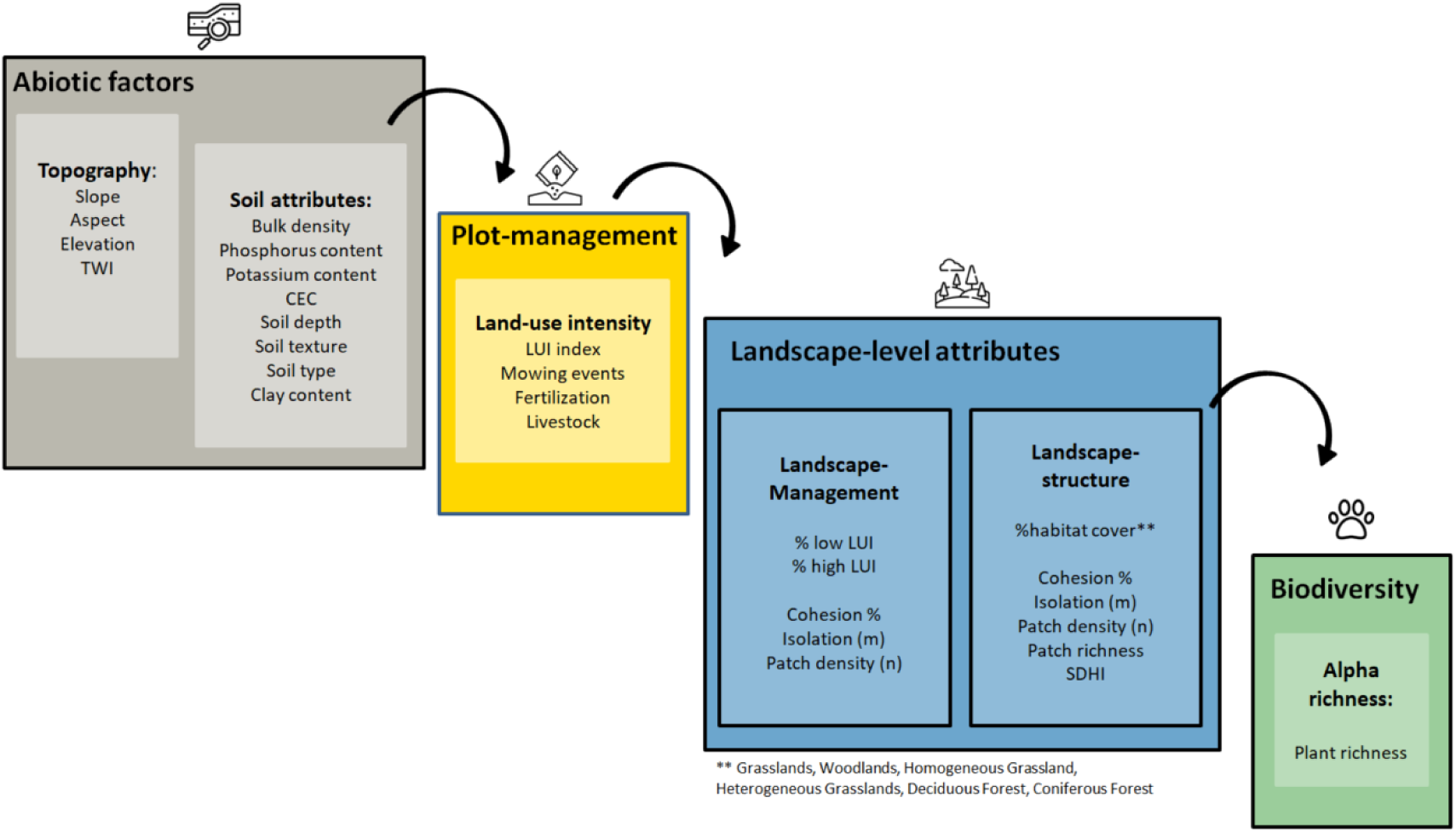
Hierarchy of effects and respective drivers within hierarchical levels tested in our hierarchy of controls approach driving multidiversity above-ground and groundwater recharge.

**Table S2.** Final scaled model coefficients of the final upscaling models for Biodiversity conservation and groundwater recharge. These models were selected using a hierarchy of controls approach and multiple likelihood ratio deletion tests (see Tutorial for the complete workflow).

| **Biodiversity conservation** | slope | p |
| --- | --- | --- |
| Intercept | 0.312 (0.013) | <0.001 |
| Slope | 0.04 (0.014) | 0.006 |
| Phosphorus content | -0.048 (0.013) | 0.001 |
| Soil depth | 0.050 (0.015) | 0.002 |
| Landscape heterogeneity | 0.044 (0.013) | 0.002 |
| **Groundwater recharge** |  |  |
| Intercept | 638.95 (9.20) | <0.001 |
| Soil type (Ren-Brau) | -62.71 (20.58) | <0.01 |
| % coniferous forest | -41.65 (8.61) | <0.001 |

### Biodiversity conservation

Since the multidiversity index encompass a large number of different taxonomic groups, finding common drivers of change might be challenging. In our model approach we detect effects acting at different levels of hierarchy, from soil and topography attributes, to landscape components.

We found that the highest contribution to the model was provided by the landscape driver related to the diversity of land-uses (SDHI) surrounding the plot at 1500m scale (explaining 15% of the variation of the model). Since we are incorporating multiple taxa in the multidiversity index, it would be expected that the SDHI had a high contribution, since higher SDHI can provide different habitats or providing feeding and nesting resources year-long contributing to higher multidiversity above-ground (Dunning *et al*. 1992). Similar results were detected in the same study region where SDHI was one of the main drivers of biodiversity aboveground (Le Provost *et al*. 2021).

Another important variable explaining 14% of model variability, was the slope of the plots. Slope, as also other topography attributes may be associated to preferred habitats or microhabitats and thermoregulation of different taxa (Davies *et al*. 2007; dos Anjos *et al*. 2022). Finally, soil attributes were also selected as important drivers, in our case, phosphorus concentration (affecting negatively biodiversity) and soil depth (showing a positive effect). Both can be associated to plant richness aboveground which will influence habitat and resource availability for a large number of taxa, especially plant diversity (Rieger *et al*. 2019). Plant richness was one of the retained drivers in our initial model, but after consultancy with experts, it was removed to avoid circularity.

### Groundwater recharge

Recharge rates are impacted by the amount and intensity of precipitation, soil and vegetation types, geology and topography (Scanlon *et al*. 2002). In geological terms, infiltration is caused by two main forces: gravity and capillarity. The rate of infiltration is determined by soil characteristics, including ease of entry, storage capacity and the saturated transmission rate through the soil. Thus, soil properties are key factor affecting groundwater recharge, which was also identified in our hierarchy of controls (explaining 26% of the variance of the final model). In the entire study region were three main classes of soil types within the range of training data in Schwäbische Alb (according to the Bundesanstalt für Geowissenschaften und Rohstoffe): Rendzina/Braunerde-Rendzina and Pararendzina, and Braunerde/Terra fusca, the latter due to the swelling and shrinking of the clay resulted in the formation of a pronounced polyhedral aggregate structure with a relatively high water permeability if not completely swollen and thus detain higher infiltration rates. In addition, it is likely that the soil type Braunerde/Terra fusca has a higher storage capacity which keeps the soil moist for longer, and consequently, higher water conductivity.

Groundwater recharge is also influenced by land-use types at the landscape scale (Scanlon *et al*. 2002). In our hierarchy of controls approach the amount of coniferous forest at the landscape scale (2000m around the plots) showed to be the most important landscape variable influencing groundwater recharge. In general, coniferous forest absorb (i.e., via interception evaporation and transpiration) more water than deciduous forests or grasslands (thus, more coniferous forest, lower groundwater recharge); the severity of reduction in recharge is influenced by the plantation density of the coniferous trees as well (Bellot *et al*. 1999; Adane & Gates 2015). Land use intensity at the landscape scale (low land use intensity) was also selected as one important driver in our final model. However, the retention of this driver got the model fit worst and increased the error associated to predictions. Thus, in order to maintain reliable predictions, we decided to leave this driver out.

## Data & Code

Data and R code to run the pipeline and reproduce the results are in my Institutional dropbox only for reviewer access - Once the paper is published, both code and data will be properly storage in a public repository. It can be find in the following link:

https://www.dropbox.com/scl/fi/inizxav8zjaar3chj5jqp/Data-UpscalingFrameworkBIORXIV.zip?rlkey=7nurebn9181qajp10l2797pg6&st=knw8iqqo&dl=0

Tutorial 1 – R script to calculate muldiversity and steps described in the paper to upscale biodiversity conservation

Tutorial 2 – R scrip to perform the steps described in the paper to upscale groundwater recharge

